# Structure of SARS-CoV-2 2′-*O*-methyltransferase heterodimer with RNA Cap analog and sulfates bound reveals new strategies for structure-based inhibitor design

**DOI:** 10.1101/2020.08.03.234716

**Authors:** Monica Rosas-Lemus, George Minasov, Ludmilla Shuvalova, Nicole Inniss, Olga Kiryukhina, Joseph Brunzelle, Karla J. F. Satchell

## Abstract

There are currently no antiviral therapies specific against SARS-CoV-2, the virus responsible for the global pandemic disease COVID-19. To facilitate structure-based drug design, we conducted an X-ray crystallographic study of the nsp16/nsp10 2′-*O*-methyltransferase complex that methylates Cap-0 viral mRNAs to improve viral protein translation and to avoid host immune detection. Heterodimer structures are determined with the methyl donor *S*-adenosylmethionine (SAM), the reaction product *S*-adenosylhomocysteine (SAH) or the SAH analog sinefungin (SFG). Furthermore, structures of nsp16/nsp10 with the methylated Cap-0 analog (m^7^GpppA) and SAM or SAH bound were obtained. Comparative analysis revealed flexible loops in open and closed conformations at the m^7^GpppA binding pocket. Bound sulfates in several structures suggested the location of the phosphates in the ribonucleotide binding groove. Additional nucleotide binding sites were found on the face of the protein opposite the active site. These various sites and the conserved dimer interface could be exploited for development of antiviral inhibitors.

## Introduction

On December 31^st^, 2019, the World Health Organization (WHO) was alerted of a pneumonia outbreak with an unknown etiology, originating in the Chinese province of Wuhan, Hubei. The etiological agent was identified as a coronavirus, closely related to the virus responsible for Severe Acute Respiratory Syndrome (SARS). The new SARS coronavirus-2 (SARS-CoV-2) causes the severe respiratory infection, Coronavirus Disease 2019 (COVID-19). Within four months, SARS-CoV-2 rapidly spread, sparking a global pandemic. The COVID-19 pandemic has also forced governments to enact “stay-at-home” orders around the world, seriously damaging the global economy (1). According to the World Health Organization, over 17 million SARS-CoV-2 infections have been confirmed, of which more than 685,000 were fatal as of Aug 1, 2020 (www.who.int). These data are similar to Johns Hopkins University tracking system (2).

The coronaviridae family of viruses causes disease in birds and mammals, including bats, camels, pigs, and humans. In lower vertebrates, pathogenic coronaviruses cause acute and severe gastrointestinal infections, fevers and organ failure. Three of the seven human-tropic coronaviruses, hCoV-229E, hCoV-NL63, hCoVB-OC43 cause only asymptomatic or mild infections, including the common cold (3). Four other human coronaviruses are linked to severe infections; including, hCoV-HKU1, a common cause of pneumonia, SARS-CoV with a 10% mortality rate, Middle East Respiratory Syndrome Virus (MERS-CoV) with a 37% mortality rate, and SARS-CoV-2 currently with a 5% mortality rate (3, 4). Among these, SARS-CoV-2 stands as the one with higher transmissibility, making its containment very difficult (5). As SARS-CoV-2 continues to spread, the need for effective vaccines and therapeutics increases. Therefore, it is urgent to study SARS-CoV-2 mechanisms of infection and replication in order to find effective targets for drug and vaccine development.

Coronaviruses have a large (∼30 kb) single-stranded, positive RNA genome that is 5′-capped, and contains a 3′-poly-A tail. The *orf1a* and *orf1ab* are directly translated, while the rest of the genome serves as template to generate sub-genomic messenger RNAs (mRNAs) transcribed from the 3’-end, which are later capped and translated (6-8). The first open reading frame produces the large non-structural polyprotein 1a (pp1a) and a programmed -1 ribosomal frameshift results in translation of the larger non-structural polyprotein 1ab (pp1ab). These polyproteins are subsequently processed into sixteen non-structural proteins (nsp1-16) that assemble to form the Replication-Transcription Complex (RTC) or function as accessory proteins necessary for viral replication (8, 9).

The components of the RTC include enzymes that regulate mRNA and genomic RNA synthesis, proofreading, and mRNA maturation. Two of these enzymes are critical for capping viral mRNAs, a tactic employed by multiple RNA viruses to avoid immune detection (10). In eukaryotic cells, mRNA capping is initiated by an RNA triphosphatase (TPase), which removes the γ-phosphate from the 5′-end of the nascent mRNA transcript, generating a diphosphate 5′-ppN end. An RNA guanylyltransferase (GTase) subsequently catalyzes the hydrolysis of pyrophosphate (PPi) from a guanidine triphosphate (GTP) molecule forming GMP, followed by the transfer of the α-phosphate of guanidine monophosphate (GMP) to the diphosphate 5′-ppN transcript end, forming the cap core structure, methylguanine-triphosphate-ribonucleotide, referred to as GpppN. The GpppN formation is followed by N^7^-methylation of the capping guanylate by a guanine-N^7^-methyltransferase (N^7^-MTase) to generate the irreversible Cap-0. Further methylation at the ribose 2′-*O* position of first nucleotide of the RNA is catalyzed by a ribose 2′-*O*-methyltransferases (2′-*O*-MTase) to generate Cap-1 and sometimes at the second nucleotide to generate Cap-2. Both the N^7^-MTase and 2′-*O*-MTase use S-adenosyl-L-methionine (SAM) as the methyl group donor (4, 10).

For coronavirus mRNA maturation, the TPase activity is mediated by nsp13, (6, 11-13) and a still elusive GTase is used to guanylate the 5’-end of the nascent mRNA. The viral non-structural protein 14 (nsp14) N^7^-MTase activity then generates the Cap-0 (14). Nsp14 is a bifunctional enzyme with independent N^7^-MTase and exonuclease domains (15). The association of nsp14 with viral non-structural protein 10 (nsp10) specifically stimulates nsp14 exonuclease activity, but has no effect on the N^7^-MTase activity (16). The coronavirus mRNAs are further modified to have a Cap-1 by the viral 2’-O-methyltransferase (nsp16). Nsp16 is a 7-methylguanine-triphosphate-adenosine (m^7^GpppA)-specific, SAM-dependent 2’-*O*-MTase (17, 18) that is activated upon binding of nsp10 (16, 19). Nsp10 is a stable monomeric protein that can also form dodecamers (20, 21), in addition to binding to nsp14 and nsp16 (16, 22). Although no specific enzymatic activity has been identified for nsp10, it is known that nsp10 is a zinc binding protein and can bind RNA (14, 20, 23). It has also been found that nsp10 could interact with human adaptor protein complex 2 (24). However, the main known function of it is the stabilization of the SAM binding pocket in nsp16 and nsp14 (19). The nsp16/nsp10-mediated 2’-*O*-methylation of coronavirus RNA is essential for preventing recognition by the host to evade immune responses that are triggered by viral mRNAs (17).

Structures of the complex of nsp16 with nsp10 have been determined for SARS-CoV and MERS-CoV (14, 18, 25, 26) and the analysis elucidated the structural basis for substrate binding and the proposed S_N_2-mechanism of methyl transfer. In order to facilitate structure-based inhibitor design, we undertook a comprehensive analysis of the structure of the SARS-CoV-2 nsp16/nsp10 heterodimer. Previously, we reported the structure in complex with the methyl donor SAM in part to make public our methods to benefit the research community (27). We herein update and extend the initial findings with a more comprehensive study. In addition to the prior reported structure with SAM bound (27), we present also the nsp16/nsp10 heterodimer complex with the product of the reaction S-adenosylhomocysteine (SAH) and the SAH analog inhibitor sinefungin (SFG). In addition, we describe the first publicly available SARS-CoV-2 structures of nsp16/nsp10 in complex with the Cap-0 analog m^7^GpppA, which facilitates detailed analysis of the changes in the conformation of flexible loops of nsp16 upon substrate binding. Furthermore, we report the only crystal structures with sulfate ions bound to the proposed RNA binding groove as well as several additional nucleotide and sugar binding sites located away from the active site. The nsp16 protein is one of the most conserved proteins of SARS-CoV-2 and related viruses and thus these high resolutions structures are expected to be useful as models for developing new antiviral therapeutics to treat COVID-19 and other diseases caused by coronaviruses.

## Results

### Two crystal form of the SARS-CoV-2 2’-*O*-MTase

The SARS-CoV-2 proteins nsp10 and nsp16 are encoded by the polycistronic *orf1ab* of the (+) ssRNA (Fig. 1A) and are released from the polyproteins by the nsp5 protease (28). Protein nsp10 is a 14.8 kDa protein released from pp1a and pp1ab, while nsp16 is a 33.3 kDa protein only created after a -1 ribosome shift that produces pp1ab polyprotein (28) (Fig. 1A). This first structure of the nsp16/nsp10 complex from SARS-CoV-2 was determined at 1.8 Å (PDB code 6WH4,((27) Fig. 1B). This crystal belonged to the space group P3_1_21 with two polypeptide chains in the asymmetric unit, with chain A (nsp16) and chain B (nsp10) forming a heterodimer. We refer to this crystal form as the “small unit cell”.

**Fig. 1.**
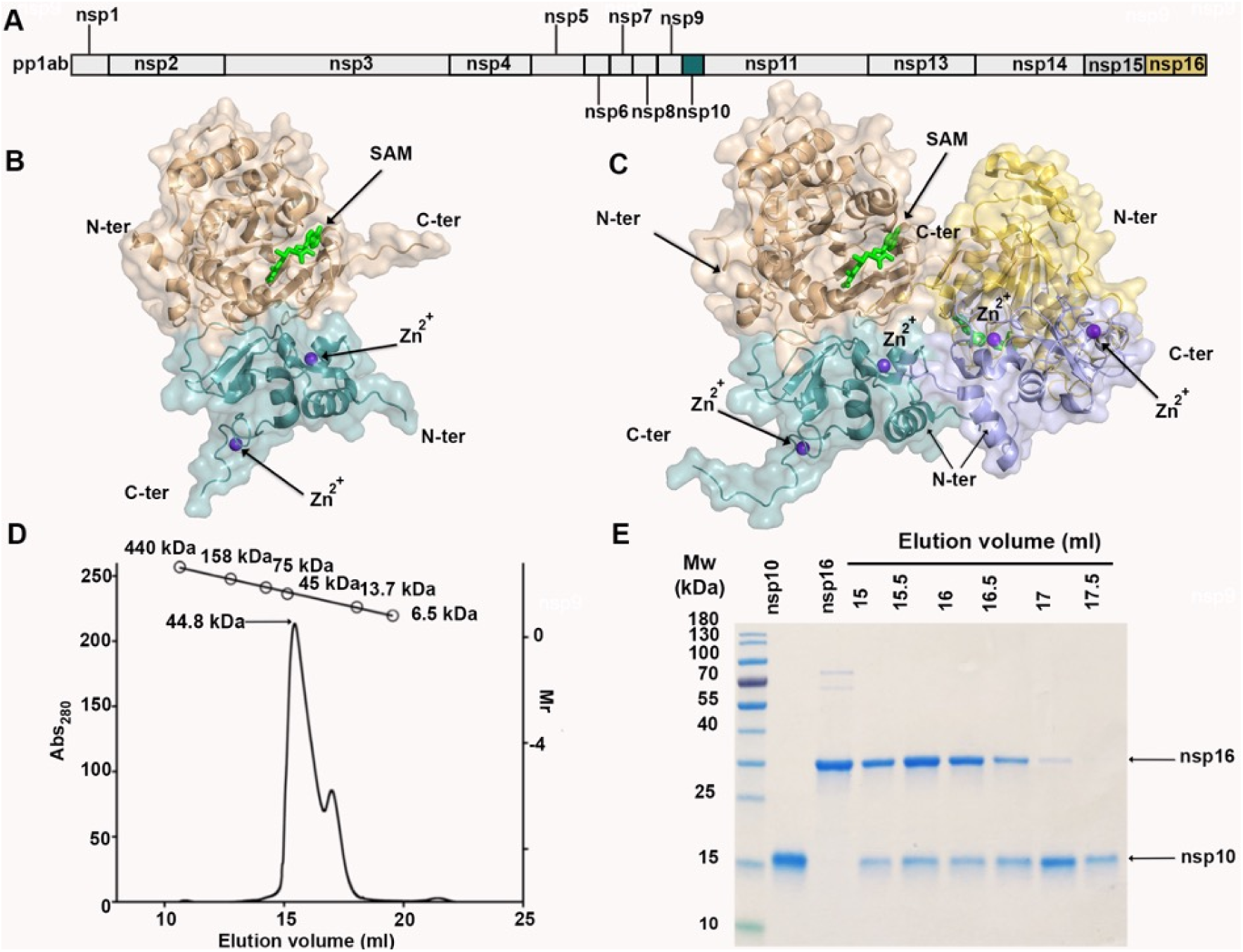
Overall structure of the nsp16/nsp10 oligomers. **(A)** Linear schematic of the *orf1a/orf1ab* protein product pp1ab prior to proteolytic processing. **(B)** Cartoon representation of the nsp16/nsp10 heterodimer of the small unit cell crystal form (PDB code 6W4H). **(C)** Cartoon representation of the two nsp16/nsp10 heterodimers in the asymmetric unit of the large unit cell crystal form (PDB code 6W75). In both panels, nsp16 is shades of tan/yellow and nsp10 is shades of teal/purple. Ligands are represented as sticks with SAM in lime green, and Zn^2+^ in purple. **(D)** Elution profile for analytical SEC with relative molecular weight plot of HMW and LMW standards shown at the top. **(E)** 4-18% SDS-PAGE gel of the elution fractions stained with Coomassie blue.

A second crystal form of the nsp16/nsp10 complex with SAM bound yielded a structure determined at 1.95 Å (PDB code 6W75). This crystal form belongs to the P3_2_21 space group and has four chains in the asymmetric unit. The four chains were arranged as a dimer of dimers with a butterfly-like shape (Fig. 1C). The two heterodimers interacted by the C-terminus of nsp16 and as well as the N-terminal of nsp10. We refer to this crystal form as the large unit cell.

To determine which stoichiometry existed in solution, we performed analytical size exclusion chromatography (SEC). In the elution profile, we observed a prominent elution peak at 15 ml that corresponded to a molecular weight (m.w.) of 45 kDa, which is close to the estimated m.w. of the heterodimer (49.8 kDa). We also observed a small peak at ∼17.5 ml, containing mostly nsp10 (Fig. 1D, E). No peak was detected corresponding to ∼90 kDa that would be consistent with four chains in a complex in solution, showing that the dimer of dimers is the result of crystal packing and that the heterodimer is the most soluble and stable form of the 2-*O*-MTase complex.

The overall structure of nsp16/nsp10 in the large unit cell was almost identical to the small unit cell structure, including the bound ligands SAM and Zn^2+^. In order to corroborate the degree of structural identity, the chains of both crystal forms were aligned using the FATCAT server (29). Alignment of nsp16 from the large unit cell (chains A and C) with the small unit cell (chain A) showed significant similarity with a raw root-mean-square deviation (r.m.s.d.) of 0.37 Å and 0.42 Å, respectively. The introduction of a flexibility factor in the alignment showed an optimized r.m.s.d. of 0.38 Å for chain A and 0.77 Å for chain C, demonstrating that the nsp16 structures in these two crystal forms are significantly similar, but that they have flexible regions. One of these flexible regions was a part of the loop formed by the residues Asp6931-Phe6947, which was disordered from the residues Lys6933 to Lys6939 in chain C, a likely explanation for the structural differences. The nsp10 alignment had a raw r.m.s.d. of 0.23Å for chain B and 0.34 Å for chain D and there were no gaps in the alignment. The optimized r.m.s.d. was 0.28 Å and 0.35 Å, respectively, indicating very low differences of these chains. Thus, both crystallographic forms were almost identical with small discrepancies caused by different conformations in the flexible loops of nsp16.

### Topology of nsp16 and nsp10 and the heterodimer interface

For this report, the topology of the nsp16 and nsp10 structures were analyzed in greater detail than previously reported (27). The nsp16 protein consist of pp1ab residues 6799-7096 and additional Ser-Asp-Ala residues at the N-terminus derived from the tobacco etch virus (20) protease cleavage site. The 2′-*O*-MTase catalytic core was comprised of a Rossmann-like β-sheet fold with the canonical 3-2-1-4-5-7-6 arrangement, in which β7 is the only antiparallel strand (Fig. 2A). This β-sheet was sandwiched by eleven α-helices and 20 loops (Fig. 2B).

**Fig. 2.**
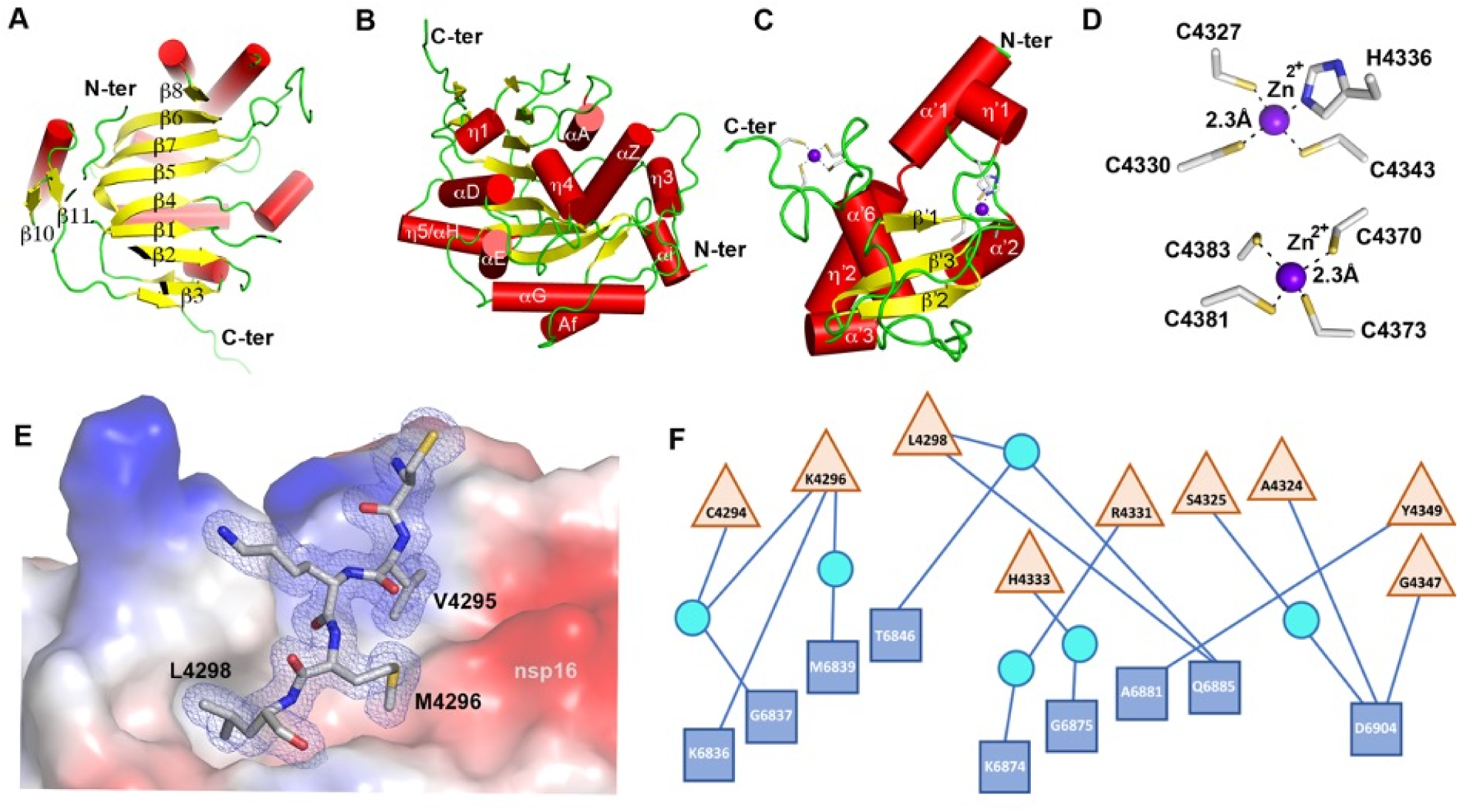
Detailed representation of nsp16, nsp10, and the heterodimer interface. **(A-C)** Cartoon representations of two views of nsp16 featuring the canonical β-sheet from nsp16 **(A)** and secondary structure of nsp16 (B) and nsp10 **(C)**. For panels A-C, α-helices are shown as red cylinders, β-strands as yellow arrows, loops as lime green strands, and zinc ions as purple spheres. **(D)** Close-up view of the two Zn^2+^ binding sites in nsp10. **(E)** Peptide fragment from nsp10 Cys4294-Leu4298 (sequence CVKML, grey sticks) interaction with the hydrophobic surface of nsp16 (shown as surface charge). **(F)** Schematic representation of residues from nsp16 (blue squares) and nsp10 (tan triangles) that interact through hydrogen bonds, represented as lines. Some interactions are mediated by water (cyan circles). For panels A-D, structural representations are based on the structure of complex with Cap (PDB code 6WVN). E and F are based on the structure of complex with SAM (PDB code 6W4H).

The nsp10 protein consisted of residues 4272-4392 of pp1a and it is composed of three β-strands (β’1, β’2, β’3), which form a central anti-parallel β-sheet. At one side of the β-sheet is the large loop that directly interacts with nsp16 and stabilizes the heterodimer complex. At the other side of this β-sheet there are six helices and loops that form two zinc fingers (Fig. 2C). In other coronaviruses, these zinc fingers are involved in non-specific binding of RNA (14, 23). The Zn-binding site 1 was coordinated by the residues Cys4327, Cys4330, Cys4336 and His4343. The Zn-binding site 2 is coordinated by Cys4370, Cys4373, Cys4381, and Cys4383 (Fig. 2D).

The residues that form the nsp16/nsp10 heterodimer interface can be divided into clusters. The clusters for nsp16 are defined as A (residues 6835-6846), B (6874-6889), C (6900-6908) and D (7042-7046). For nsp10 they are defined as cluster I (4293-4300), II (4322-4337) and III (4346-4349) (14, 18). Almost all of the interface contacts between nsp16 and nsp10 are formed by hydrophobic interactions between nsp10, cluster I (Val4295, Met4297, Leu4298) and cluster A (Pro6835, Ile6838, Met6839, Val6842, Ala6843), cluster B (Val6876, Pro6878, Ala6881) and cluster D (Leu7042, Met7045) of nsp16 (Fig. 2E,). The remaining interactions at the interface are mediated by hydrogen bonds and these hydrophilic interactions consist of five direct contacts between residues Lys4296, Leu4298, Ala4324, Tyr4349, and Gly4347 of nsp10 with Lys6836, Gln6875, Ala6881, and Asp6904 of nsp16, plus eight water-mediated interactions (Fig. 2F).

### The binding of SAM and SAH and the pan-MTase inhibitor SFG to the methyl donor binding site

The nsp16 protein catalyzes the transfer of the SAM methyl group to the Cap-0 structure, generating the reaction products SAH and Cap-1. This reaction can be inhibited by SFG, a 5′-aminoalkyl analog of SAH used as a pan-inhibitor of methyltransferases (Fig. 3A). In order to identify potential structural differences between SAM, SAH, and SFG in the SAM binding cleft, we also determined the structures of nsp16/nsp10 in complex with SAH (PDB code 6WKQ) and SFG (6WJT) at 2.0 and 1.98 Å resolution, respectively. These structures showed that SAM binds to a negatively charged cleft formed by αA, αz, αD the loops L5, L9, L11, in nsp16 (Fig. 3B). The adenosine moiety is stabilized by residues Phe6947, Asp6912, Leu6898, Cys6913 and Met6928. The sugar moiety is stabilized by the residues Gly6871, Asp6897 and by two molecules of water that interact with Asn6899. The methionine moiety interacts with Asp6928, Tyr6845, Asn6841 and Gly6871. Notably, SAH and SFG interact with the same residues as SAM without significant modifications in the site or the overall structure (Fig. 3C).

**Fig. 3.**
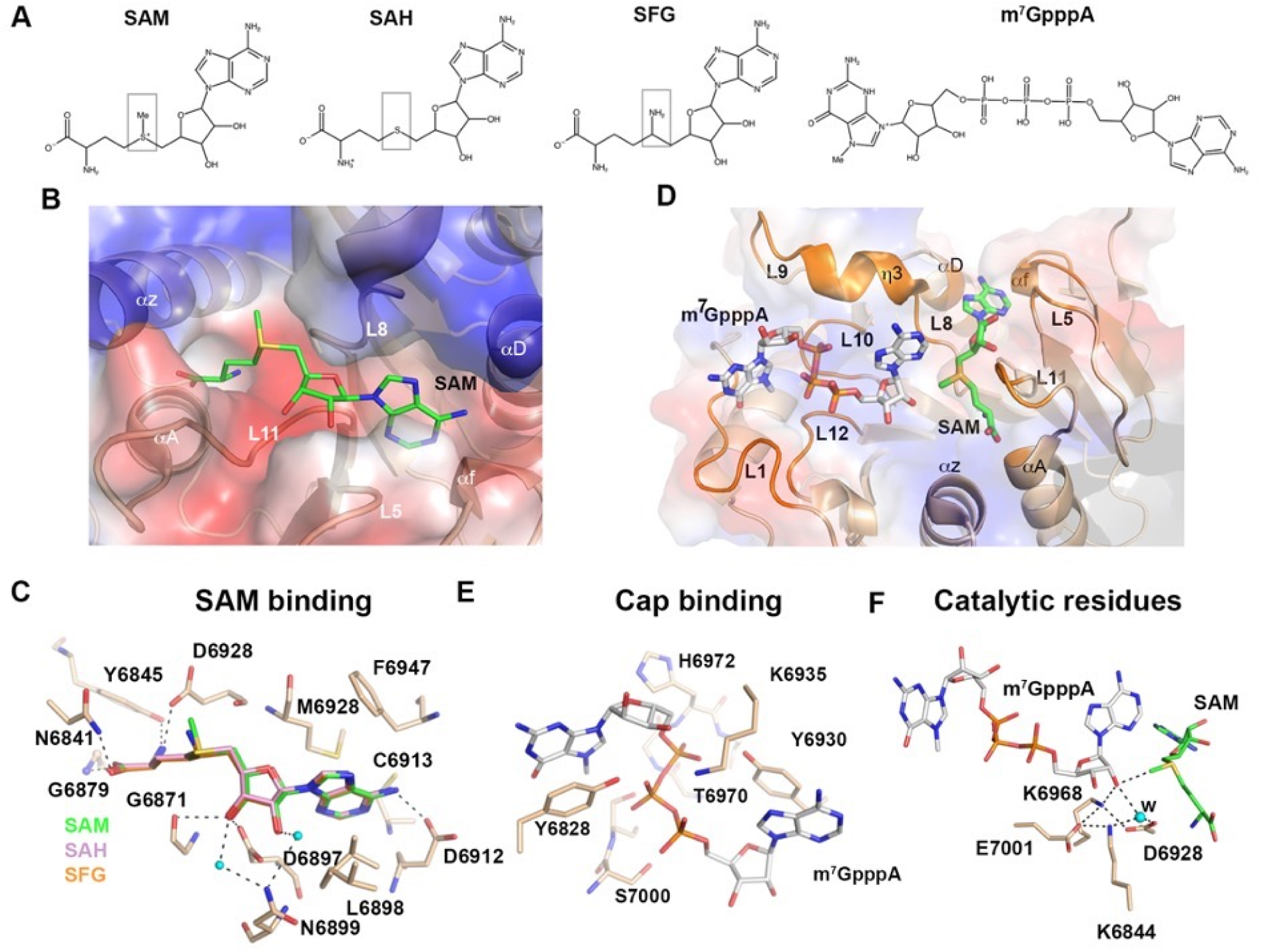
Ligand interactions and catalytic site. **(A)** Chemical structures of the methyl donor SAM, the product after methyl transfer SAH, and the SAH analog inhibitor SFG and the Cap-0 m^7^GpppA. Boxes highlight differences in the chemical structures compared with SAM. **(B)** Cartoon and surface charge representations of the nsp16 SAM binding cleft occupied by the SAM (green sticks). **(C)** Close-up view of overlay of nsp16 structures with the ligands SAM (lime green, PDB code 6W75), SAH (pink PDB code 6WJT) and SFG (PDB code 6WQK). **(D)** Cartoon and surface charge representations of nsp16 (PDB code 6WVN) of the SAM binding cleft occupied by the ligand SAM (lime green sticks) and the Cap binding site occupied by m^7^GpppA (gray sticks). **(E)** Detailed view of residues that coordinate m^7^GpppA in the binding site as tan sticks. **(F)** Close-up view of the side chains of catalytic residues showing orientation of the methyl group in SAM in proximity to the acceptor 2’-OH group in m7GpppA. Dashed lines indicate the interactions between the residues in the active site and small cyan dots indicate water (w). Oxygen as red sticks, nitrogen in blue, phosphate in orange and sulfur in yellow.

### Binding of SAM and m^7^GpppA to the nsp16 catalytic site

In addition to the methyl donor SAM, the nsp16 catalytic reaction requires a Cap-0-mRNA with an adenosine in position 1 of the single stranded RNA (m^7^GpppA-RNA). The crystal structure of the ternary complex of nsp16/nsp10 with m^7^GpppA and SAM was previously described for the more distant relative MERS-CoV (26), but not for SARS-CoV, wherein the binding of the mRNA was only modeled (14). Herein, we describe the first publicly available structure of a lineage B beta-coronavirus nsp16/10 heterodimer in complex with the analog of the substrate of the methylation reaction, m^7^GpppA.

The molecule m^7^GpppA bound to the Cap High Affinity Binding Site (HBS) which is a positively charged surface on nsp16 formed by the loops L1, L8, L9, L10 and L12, αD, and the η3 (Fig. 3D). The guanosine ring of m^7^GpppA is stacked with Tyr6828. The phosphate groups are mostly stabilized by side chain atoms of Tyr6828, Tyr6930, Lys6935, Thr6970, Ser6999, and Ser7000, and by the main chain atoms of His6972 and Ser7000 of loops 10 (residues 6970-6975) and 12 (residues 6994-6997) (Fig.3E). The adenosine sugar interacts with side chain atoms of Lys6844, Lys6968 and with Asp6928 through a water molecule. The adenine moiety is stabilized by stacking interaction with side chain of Tyr6930 and it is in close proximity with the SAM binding cleft. These interactions are also found in the structure of MERS-CoV nsp16 in complex with Cap-0 (PDB code 5YNM (26)).

The high quality of the crystal structure of the nsp16/10 complex with m^7^GpppA bound facilitated detailed analysis of the catalytic site. The protein nsp16 contains the highly conserved residues Lys6839, Asp6928, Lys6968, and Glu7001, comprising the canonical catalytic motif (K-D-K-E) conserved among class I MTases (14, 18, 30). These residues are close to the SAM methyl group that is transferred to the 2′-OH on the m^7^GpppA. NZ of the Lys6968 is in close interaction with the 2′-OH of Cap-0 and possibly activates this oxygen for the nucleophilic attack of the methyl group in SAM (Fig. 3F). In the structure of nsp16/nsp10 with the m^7^GpppA and SAM bound, we detected the presence of one molecule of water (Fig. 3F) that might participate in the stabilization of intermediate catalytic states (14, 18, 31). Although the nsp16 MTase reaction was previously characterized as Mg^2+^-dependent, we did not observed this metal bound and it is likely that Mg^2+^ is involved only in transitory states of catalysis or stability of the protein as previously suggested for dengue virus 2′-*O*-MTase (32).

### The flexibility of the m^7^GpppA site in nsp16

Several studies of SARS-CoV-1 nsp16 demonstrated the stabilization of the SAM cleft upon nsp10 binding and that m^7^GpppA-RNA binding causes conformational changes (14). However, no crystal structures supporting these conformational changes have as yet been described. The HBS is surrounded by Loop 1 (residues 6824-6834) and a loop formed by L8-η-3 -L9 (residues 6930-6943). These flexible loops are visible for the first time in our structures and thus we could analyse the diverse conformations of nsp16/nsp10 complexes presented in this work.

First, we analyzed structures of the heterodimer with only SAM bound from the large unit cell form (Fig. 4A, green) with the small unit cell form (blue). There are only minor conformational differences for loop 1, however, the L8-η-3 -L9 loop shows a more “open” conformation in the large unit cell and a more “closed” conformation for the small unit cell structure when only SAM is bound. This analysis corroborated that this specific region is flexible and that these loops likely transit from an “open” to a “closed” state in absence of m^7^GpppA.

**Fig. 4.**
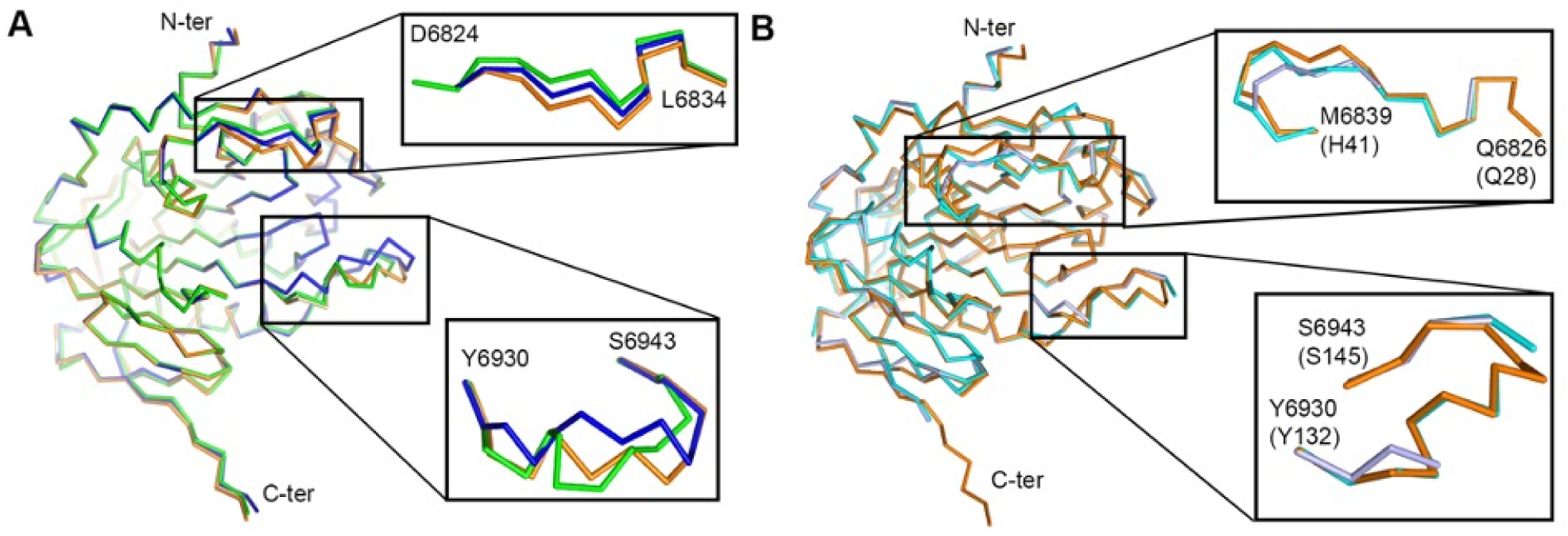
Structural alignment of nsp16 in presence and absence of m^7^GpppA. **(A)** Alignment of the C-α chain of nsp16 from SARS-CoV-2 in complex with SAM from the small unit cell (blue, PDB ID 6W4H), nsp16 with SAM from the large unit cell crystal form (green, PDB code 6W75, green) and nsp16 in complex with SAM and m^7^GpppA (orange, PDB code 6WVN). Two flexible loops are enlarged in insets. **(B)** Alignment of the C-α chain of nsp16 from SARS-CoV-2 in complex with SAM and m^7^GpppA (orange) with MERS-CoV nsp16 with SAM alone (light blue, PDB code 5YN6) or with SAM and m^7^GpppA (cyan, PDB code 5YNM), numbering of MERS structure in parenthesis.

In order to evaluate the position of loops when m^7^GpppA is bound, we analyzed the structural alignment between the high-resolution heterodimer with SAM bound from the small unit cell (Fig. 4A, blue) and the heterodimer in complex with m^7^GpppA and SAM (orange). This alignment has an r.m.s.d. of 0.55 Å and we observed that the presence of m^7^GpppA induces a stable open conformation of residues 6930-6943, which was found also in the heterodimer in complex with m^7^GpppA and SAH (r.m.s.d.=0.12 Å, not shown in Fig. 4A). The main conformational changes were observed near residues 6936-6939, which were displaced in the open conformation and appeared to mitigate clashes with the guanidine sugar and the 1^st^ phosphate group of the m^7^GpppA and Lys6935 (Fig. 5). These changes indicated that the presence of m^7^GpppA stabilized an open state of the Cap-0 binding site, which could facilitate the release of the product upon methylation, this conformation was also observed in other 2’-MTase from SARS-CoV-2 structure (PDB code 6WKS) (33).

**Fig. 5.**
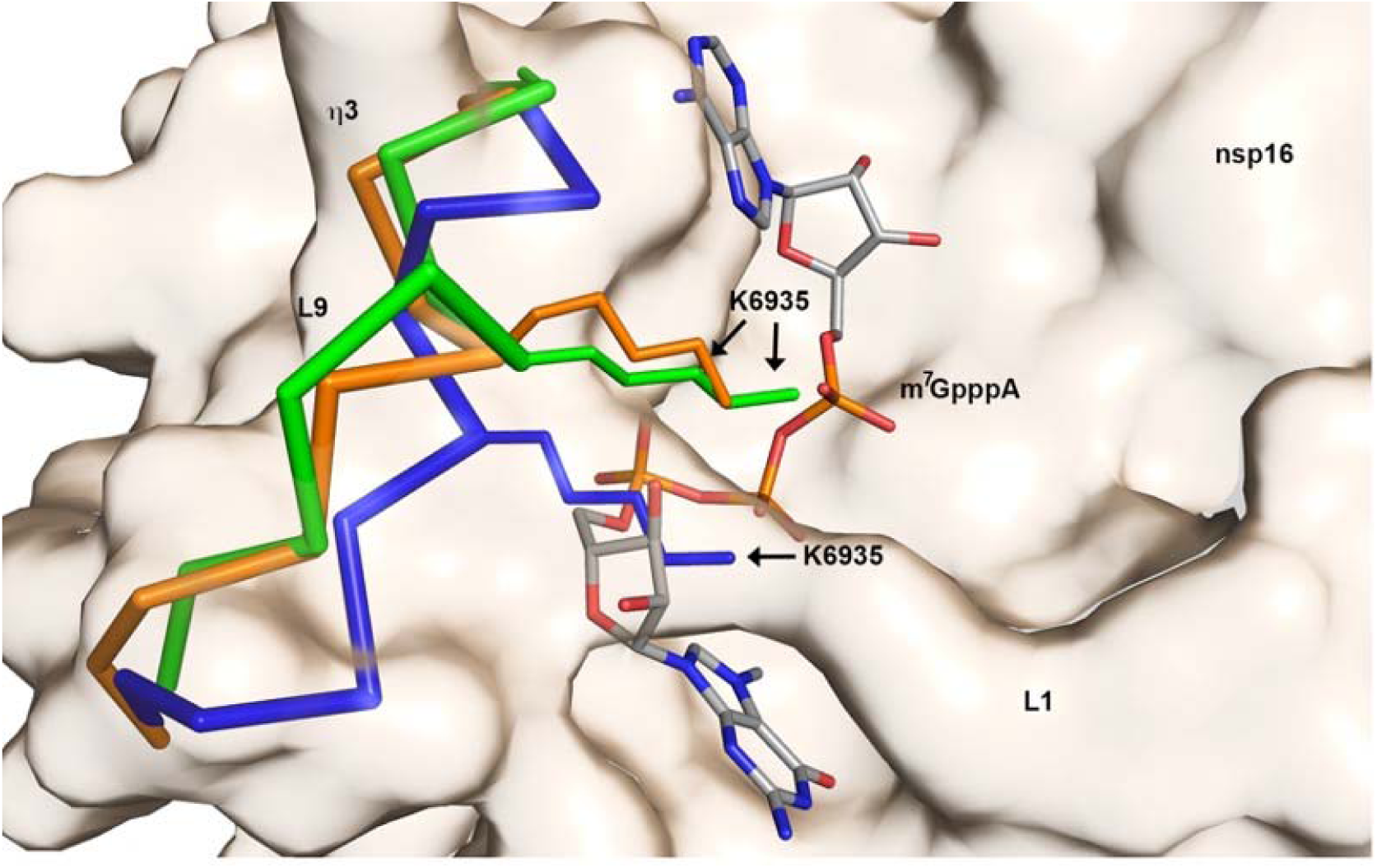
Displacement of the Lys6935 upon m^7^GpppA binding. The flexible loops L8-L9 of nsp16, large unit cell with SAM bound PDB code 6W75 (green), small unit cell SAM bound PDB code 6W4H (blue) and SAM+ m^7^GpppA, 6WVN (orange) are shown as C-α ribbon. The Lys6935 as sticks and the m^7^GpppA in gray for carbon; red, oxygen; blue nitrogen and orange phosphate. The surface shown in tan corresponds to the structure PDB code 6W4H.

Furthermore, the C-α chain of nsp16 from SARS-CoV-2 with Cap-0 and SAM bound (Fig. 4B, orange) was aligned with MERS-CoV nsp16 with SAM (violet) or with cap-0 and SAM bound (cyan) over 293 residues. In the absence of the Cap-0, the MERS-CoV nsp16 residues corresponding to residues 6934-6940 in SARS-CoV-2 are disordered, confirming that this loop is flexible. In contrast, Loop 1 and residues 6930-6943 from MERS-CoV m^7^GpppA-nsp16 complex aligns with the open conformation observed in the SARS-CoV-2 m^7^GpppA-nsp16 complex. This indicates that both 2′-*O*-MTases have the same open conformation upon Cap-0 binding.

### Sulfates align to the RNA binding groove

The 2′-*O*-MTase nsp16/nsp10 possibly binds the RNA tail in the positively charged nucleotide binding groove, also known as the low affinity binding site (LBS). There is, as yet, no structural evidence of the arrangement of this part of the viral RNA in the protein and only prediction models have thus far been published (14, 33, 34). Sulfates are known to bind to the same positions as phosphates in proteins. Since the first small unit cell structure (PDB code 6W4H) was obtained from crystallization conditions with polyethylene glycol (PEG), which did not allow for soaking with substrates, crystals were screened for suitable conditions for soaking with substrates followed by cryoprotection with 2 M lithium sulfate (see methods). We speculated that these molecules of sulfate could indicate the possible phosphates of the RNA molecule. All of the structures with m^7^GpppA also had molecules of sulfates bound at distinct sites and could be superimposed to analyze the position of sulfates (Fig. 6A-C). Sulfate 1 (S1) was in a position next to the SAM cleft in two alternative conformations, which could indicate the importance of charged molecules nearby the catalytic site (Fig. 6B). Sulfate 2 (S2) seems to mimic the phosphate group between the first and the second nucleotide in the RNA and is followed by a zig-zag line of other sulfates (S2-S5) along the positively charged LBS from nsp16 and the extension groove of nsp10. The compilation of these results suggested that the nucleotide binding groove might accommodate four to five nucleotides from the viral m^7^GpppA-RNA (Fig. 6B). Thus far, this is the only experimental and structural evidence that reveals the possible position of the RNA in the nucleotide binding groove.

**Fig. 6.**
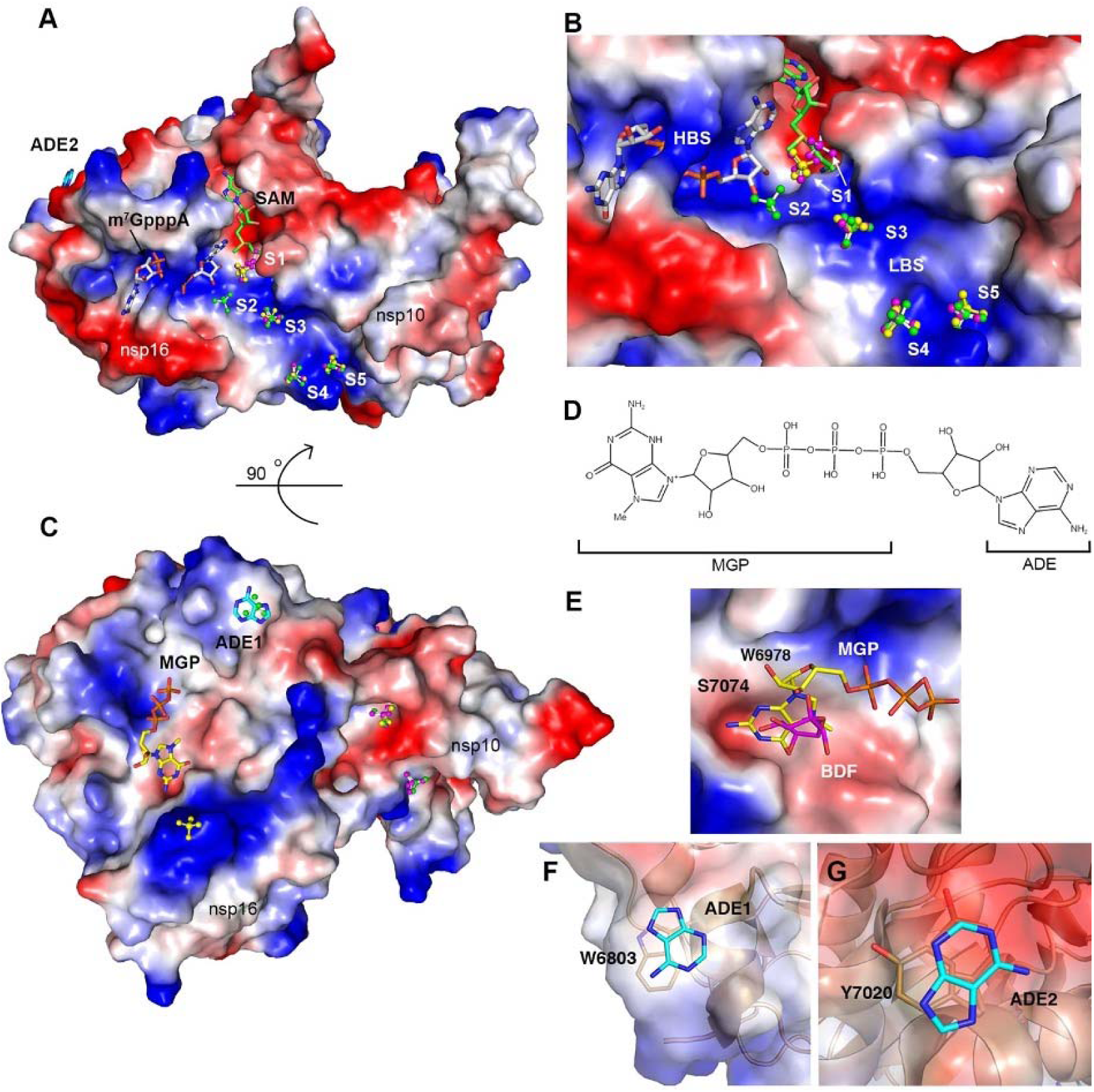
Sulfates and nucleotides binding sites of nsp16. **(A)** Surface charge representation, of nsp16/nsp10 with SAM and m^7^GpppA bound in green and gray respectively (PDB code 6WVN) and **(B)** sulfates in balls and sticks along the nucleotide binding groove numbered from the catalytic core to the nsp10 extension (S1-S5). Positive charges in blue and negative charges in red. The sulfates in the overlayed structures are designated by color according with their corresponding PDB code: 6WRZ, 6WVN (yellow), and 6WQ3 (pink). m^7^GpppA is shown as gray sticks and SAM in green sticks. HBS, high affinity binding site; LBS, low Affinity Binding site. **(C)** 90° rotation of the complex showing the secondary binding sites MGP and ADE1 along with additional sulfates. **(D)** Schematic representation of m^7^GpppA and name of the different moieties in the m^7^GpppA structure. **(E)** Surface charge representation of nsp16 MGP binding site with MGP as (yellow sticks) and BDF from structure 6W4H (pink sticks). **(F**,**G)** Cartoon and surface charge representation of the of the adenine moieties (ADE1 and ADE2). For all ligands, carbon as indicated sticks, nitrogen in blue, oxygen in red, phosphates in orange and sulfate in yellow.

#### Identification of three additional m^*7*^*GpppA binding sites in nsp16*

The present comprehensive study of the interaction of m^7^GpppA with nsp16/nsp10 resulted in the unexpected finding of nucleotides in non-catalytic sites of the structures (Fig. 6A and 6C). Although relatively short soaking times were tested to avoid non-specific binding of m^7^GpppA, nucleotides were consistently found at three different positions, additional to the active site. Here we describe for the first time three non-catalytic nucleotide binding sites as result of soaking the crystals with m^7^GpppA and SAM or SAH. One of the sites showed binding of the guanine and phosphate moiety from m^7^GpppA (designated MGP, Fig. 6D) and is located on the back surface of the protein (Fig. 6C). This site was found occupied in three different experiments. The guanine moiety interacts with the hydrophobic surface formed by Trp6987 and Leu6855 and the NH_2_ is stabilized by Ser7074 in a small negatively charged cavity (Fig. 6E). The 2′-OH from the sugar is stabilized by hydrogen bonds between a molecule of water, Asp6811, the first phosphate and the N1 from the Trp6987. The adenosine moiety of the m^7^GpppA ligand was disordered. In the structure without the cap (PDB code 6W4H), this same site was occupied by β-D-fructopyranose (BDF) (Fig. 6E) indicating that this site is not nucleotide-specific.

Another binding site (ADE1) was occupied by an adenine (ADE) moiety that likewise is derived from the m^7^GpppA (Fig. 6C and 5D). The ADE stacked with Trp6803 (Fig. 6F). The third binding site (ADE2) is also occupied by an adenine ring stacking with Tyr7020 (Fig. 6A and 6G). These non-catalytic nucleotide binding sites might represent interactions that would mimic the interaction between nsp16 and the ribonucleotides of the capped mRNA.

## Discussion

The SARS-CoV-2 pandemic has yielded a world-wide effort to understand the molecular mechanisms involved in virus transmission, virulence, and replication (1). The ultimate goal is to identify viral proteins that are amenable for drug targeting and epitopes suitable for vaccine development. Although, previous studies conducted in SARS-CoV and MERS-CoV paved the way for drug discovery and vaccine development, no approved treatments were fully developed (35). Thus, in order to ensure an accurate approach for drug discovery, we present a comprehensive study of the structure of the SARS-CoV-2 2′-*O*-MTase complex. In addition to the structures reported here, similar structures of the nsp16/nsp10 complex have subsequently been determined by other groups; including structures with SAM (PDB codes 6W61 (36), 7BQ7 (37), 7C2I, and 7C2J (38), with SFG (PDB code 6YZ1, (34)). Independently, another structure of nsp16/nsp10 in complex with m^7^GpppA + SAM was deposited (PDB code 6WKS (33)), although this structure was released only at a later date.

Previous studies determined that the 2′-*O*-MTases of SARS-CoV-1 and MERS-CoV, which share 93-99% and 59-66% identity to SARS-CoV-2 MTase, respectively, are heterodimers formed by binding of nsp10 to nsp16 (16, 22). In addition, their structures have provided some insight into nsp10 dependent activation of nps16 MTase activity and catalysis (14, 18, 26). As variation at the primary sequence level can impact both local and overall structure, ligand binding and structure-based drug design can also be affected. Thus, high resolution structures of the 2′-*O*-MTase from SARS-CoV-2 are needed to best inform drug discovery for COVID-19.

This study demonstrated that the binding site for the methyl donor SAM is highly conserved, especially the canonical Gly-X-Gly motif located at the end of the β1 and αA and Phe6949 as found in almost all class I MTases (39, 40). SAM analogs have been proposed as antimicrobials targeting MTases of fungi and parasites (41, 42). Here we determined the structure of nsp16/nsp10 with the pan-MTase inhibitor SFG bound and showed that it has nearly identical interactions with amino acids side chains as the natural substrate SAM. The high resolution of the structures with SAM, SAH, and SFG bound could facilitate computational design of the small molecules that bind into the SAM binding cleft that might have a higher specificity for SARS-CoV-2. Further, the conservation of the binding cleft residues across the beta-coronaviruses suggests that an inhibitor designed for SARS-CoV-2 could also be a broader spectrum inhibitor, which could target other coronavirus nsp16 proteins.

The Cap-0 binding site offers another position in nsp16 that could be a target for small molecule inhibition. Although the published structure of nsp16/nsp10 from SARS-CoV was elucidated with SAM bound, only a computation model of the interaction of nsp16 with m^7^GpppA-RNA was previously available (14). In order to study the different arrangements in this structure upon cap binding, crystals were soaked with m^7^GpppA in presence of both SAM and SAH. The resulting structures identified the residues that interact with the Cap-0 and showed that conformational changes in the Cap-0 binding site could occur during catalysis. Of particular concern for the potential development of a more broad-spectrum inhibitor that could target the Cap binding site are two nearby loops that are variable in sequence across the beta-coronaviruses that could have an impact on its function within the cell or catalysis (Fig. 5). Our analysis of overlapped structures of nsp16 from the complex nsp16/nsp10 SARS-CoV-2 with the structure of SARS-CoV with the Cap binding site unoccupied demonstrated that these amino acid differences do not affect the overall structure of the complex, but rather are highly flexible loops that are then stabilized upon binding of the Cap. Stabilization of these loops may be critical to obtain a high affinity small molecule inhibitor directed at this site. The high-resolution structure of SARS-CoV-2 with the cap bound should facilitate design of such a molecule.

A computational model of SARS-CoV 2′-*O*-MTase in complex with RNA was proposed previously using the structure of vaccinia virus MTase as the model (PDB code 1AV6) (14, 33, 34). This model suggested that the residue Asp^75^ in SARS CoV (Asp6873 in SARS-CoV-2), confers the selectivity of the Cap binding site for m^7^GpppA over m^7^GpppG due to steric hindrance of the Asp residue with the NH_2_ at position 2 of the guanidyl (14). This residue is conserved in SARS-CoV-2. However, in all our structures with Cap bound, the position 2 of the adenylate from m^7^GpppA is 7 Å away from the oxygen of the Asp6873, suggesting this residue is unlikely involved in the selectivity of the RNA-capped substrate in SARS-CoV-2. Indeed, recent studies show that m^7^GpppG-RNA is 2′-*O*-methylated by nsp16/nsp10 from SARS-CoV-2, but at a lower efficiency that m^7^GpppA-RNA (33, 43), indicating that this site can accommodate m^7^GpppG.

In addition to the ligand binding sites, we explored the positively charged nucleotide groove, also named LBS, which leads from the catalytic core toward nsp10. Several efforts to obtain a short RNA bound into this groove of crystals have thus far been unsuccessful, although several computational models have been recently published and are already available for SARS-CoV (14) and SARS-CoV-2 (14, 33, 34). However, no structural evidence of RNA binding was reported. Thus, in order to obtain experimental evidence of the possible accommodation of the phosphate groups for four to five ribonucleotides of the mRNA that will directly interact with nsp16/nsp10 groove, the crystals were cryoprotected with lithium sulfate. The unique arrangement of sulfate molecules in the LBS provides experimental evidence of the possible accommodation of the phosphate groups for four to five ribonucleotides of the mRNA that will directly interact with nsp16/nsp10 groove. We speculate that small charged molecules could be designed to prevent binding of mRNA, which could impair the efficiency of the MTase reaction. The potential advantage of targeting a site away from the SAM or Cap binding sites could avoid cross-inhibition of human MTases and preventing toxicity.

To this end, this study revealed previously unnoted features in the structure that could be advantageous for the design of new therapeutics. First, a possible second nucleotide binding site occupied by a guanine moiety was identified on the back of surface of nsp16 when the crystal was soaked with m^7^GpppA. This site is not specific for guanine, since we observed also BDF in the structure when cryoprotected with sucrose in the absence of m^7^GpppA. The other sites bound the adenine moiety of m^7^GpppA. These sites could be positions for interaction with the mRNA, but also have been postulated as allosteric sites (33). The binding of adenine at these positions via stacking with Tyr7020 or Trp6803 suggesting other bases could also bind at these sites indicating that if they are allosteric sites, the ligand may not be highly specific. Thus, these binding sites need to be further studied in order to determine whether the binding of other molecules might affect the activity of the enzyme, and if these newly defined binding sites could be used as potential new candidates for developing inhibitors.

A final focus for development of inhibitors against nsp16 is to target the interface with its activator nsp10. Derived nsp10-peptides were developed to target SARS-CoV-1 nsp10 that interfered with nsp10 binding to nsp16 resulting in inhibition of its activity (44). In our analysis, we found that the residues that form the interface between nsp16 and nsp10 are 100% conserved with SARS-CoV-1. Thus, we predict that small molecules that target the nsp16/nsp10 interface as well as specific small peptides could also be highly effective inhibitors that would have broader spectrum activity. An advantage of an inhibitor that binds specifically to nsp10 could even have broader implications and potentially also inhibit the exonuclease activity of the N^7^-MTase nsp14, which likewise uses nsp10 as its activator (16, 45).

One problem for the development of antiviral compounds against SARS-CoV-2 is the potential impact of emerged mutations in its genome. The new emerging mutations on SARS-CoV-2 have recently been mapped on the structures of the proteins (https://coronavirus3d.org) (7). Few mutations have emerged for nsp10 and none impact structure. Interestingly, some emerging mutations have been identified for nsp16, but none of these are predicted to disrupt the structure of nsp16. Further, review of these data showed that no mutations of residues at the interface of nsp16 and nsp10 have as yet been detected in SARS-CoV-2 viral variants isolated from around the globe (7). Thus, it is reasoned that a mutation affecting nsp16 activity could result in early immune detection followed by more rapid clearance of virus. In support of this, recent studies have suggested that stimulation of an interferon response early in SARS-CoV-2 infection could result in less severe disease, particularly in younger individuals (46). In addition, previous reports for Mouse Hepatitis Virus (MHV) showed that mice treated with a 29 amino acid peptide derived from the loops of nsp10 that interact with nsp16 inhibit the MTase activity of nsp16. This peptide also promoted survival of the treated mice from the infection and protection was mediated by the interferon response (44). Altogether, this analysis suggests that immunological studies of the impact of nsp16 inhibition could benefit from identification of an inhibitor that can be employed for study of the virus in vitro or in animals. Such an inhibitor could also be used during early infection to stimulate immunity, reducing the likelihood for development of severe disease. The structural work found in this study will help with these next stages in our understanding of MTase modification of mRNA and improved treatments for COVID-19.

## Materials and Methods

### Protein purification and analytical SEC

All methods for cloning, expression and purification of nsp16 and nsp10 were previously detailed (27). In order to confirm that the nsp16/nsp10 complex was formed, we performed analytical Size Exclusion Chromatography (SEC) using a Superdex 200 10/30 column with 10 mM Tris-HCl, pH 7.5, 150 NaCl, 2 mM MgCl_2_, 1 mM TCEP and 5% glycerol. The standard calibration curve was obtained using combined low molecular weight (LMW) and high molecular weight (HMW) Gel filtration Calibration kits (GE Healthcare). The resulting peaks from the elution of the protein were fractionated in 0.5 ml. Each fraction was collected and 8 μl of sample was denatured with Laemmli buffer, then separated using 4-15% gradient SDS-PAGE (Bio-Rad).

### Crystallization, data collection, and structure refinement

The nsp16/nsp10 complex + SAM was set up as 2 µl crystallization drops (1µl protein:1µl reservoir solution) in 96-well plates and equilibrated using commercially available Classics II, PEG’s II, AmSO_4_, Anions and ComPAS Suites (Qiagen). Diffraction quality crystals appeared after 5-10 days in 78 conditions, 118 crystals of various complexes were frozen, and 57 data sets were collected. The crystals were soaked, cryoprotected and flash frozen for data collection as follows.

The small unit cell crystal (6W4H) was cryoprotected with 25% of sucrose in the well solution and the large unit cell crystal (6W75) – with 4M sodium formate (Table 1). In order to obtain complexes with SAH and SFG, crystals were transferred into the 10 µl drops with their well solutions supplemented with 5mM of SAM or SFG, soaked for 3 h, cryoprotected with 4M sodium formate or 2M LiSO_4_ and flash frozen. In the attempt to observe the complexes of nsp16/nsp10 with SAM + m^7^GpppA and SAH + m^7^GpppA crystals were transferred into 10 µl drops containing 5 mM SAM or SAH and 0.5 mM of m^7^GpppA in their respective well solutions, soaked for various amount of time from 3 min to 6 hrs and flash frozen. 2 M LiSO_4_ was used as a cryoprotectant cryoprotectant in an attempt to observe the binding of sulfates on the places of phosphates from the RNA in the RNA binding groove. Crystals grown in polyethylene glycol (PEG) conditions were found not suitable for these soaks due to a phase separation of m^7^GpppA in presence of PEG.

**Table 1.**
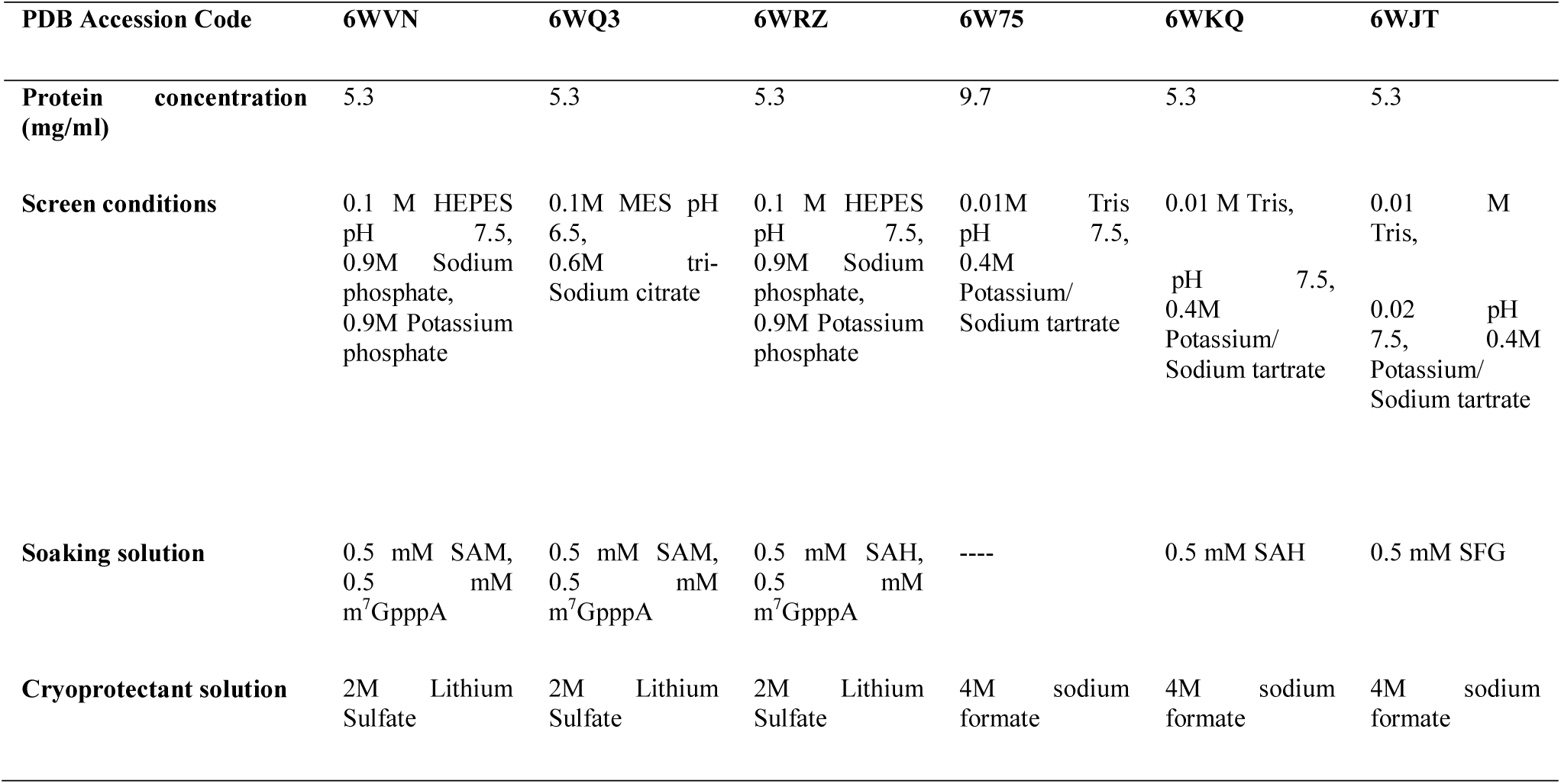
Crystallization, soaking and cryoprotection conditions.

### Data collection and processing

Almost 120 crystals were screened, and 57 data sets were collected at the Life Sciences-Collaborative Access Team (LS-CAT) beamlines D, G and F at the Advanced Photon Source (APS) at the Argonne National Laboratory. All the data sets reported here were collected at the beamline F. Images were indexed, integrated and scaled using HKL-3000 (47). Seven structures were chosen to be described in this manuscript (Table 2).

**Table 2.**
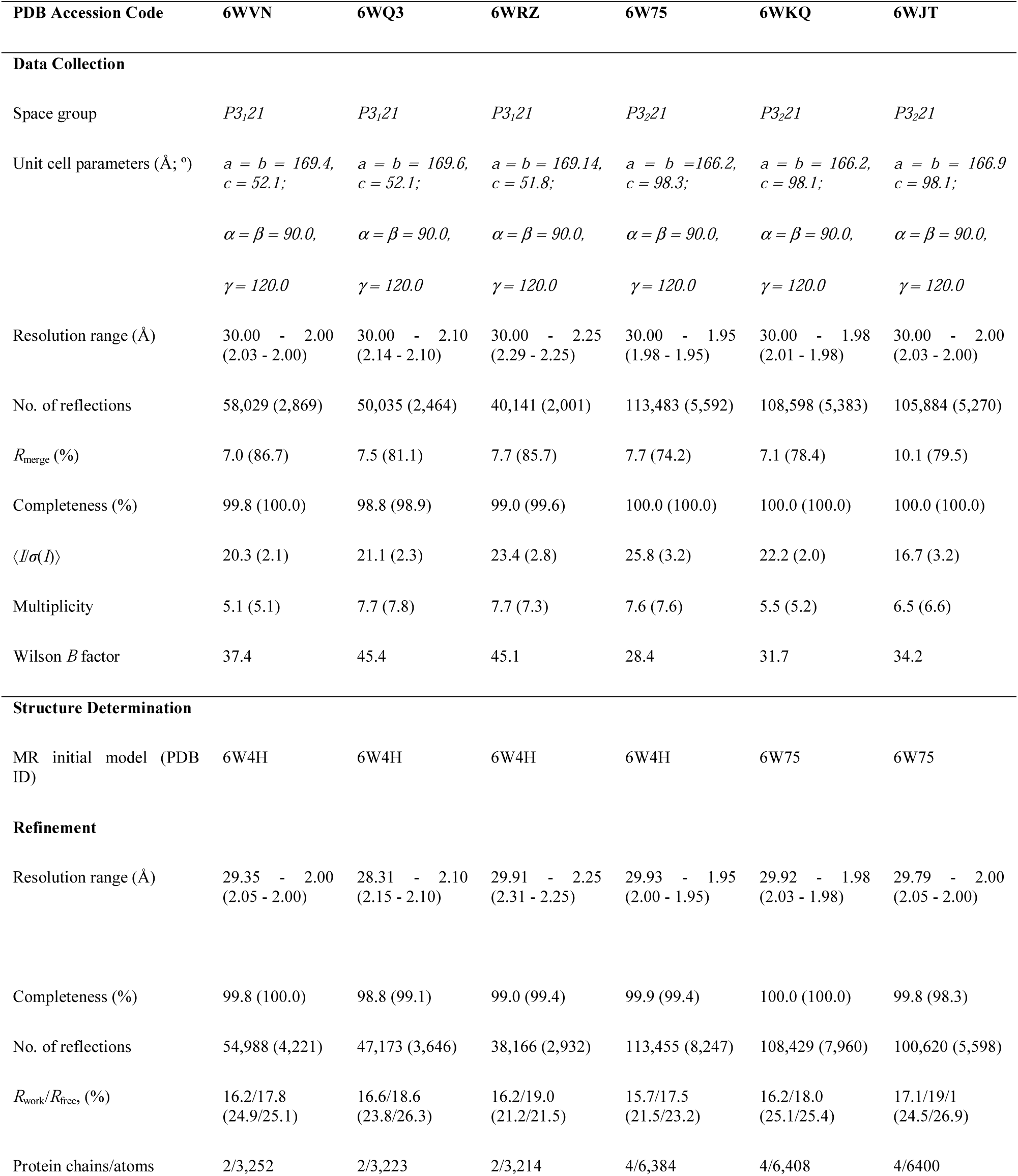
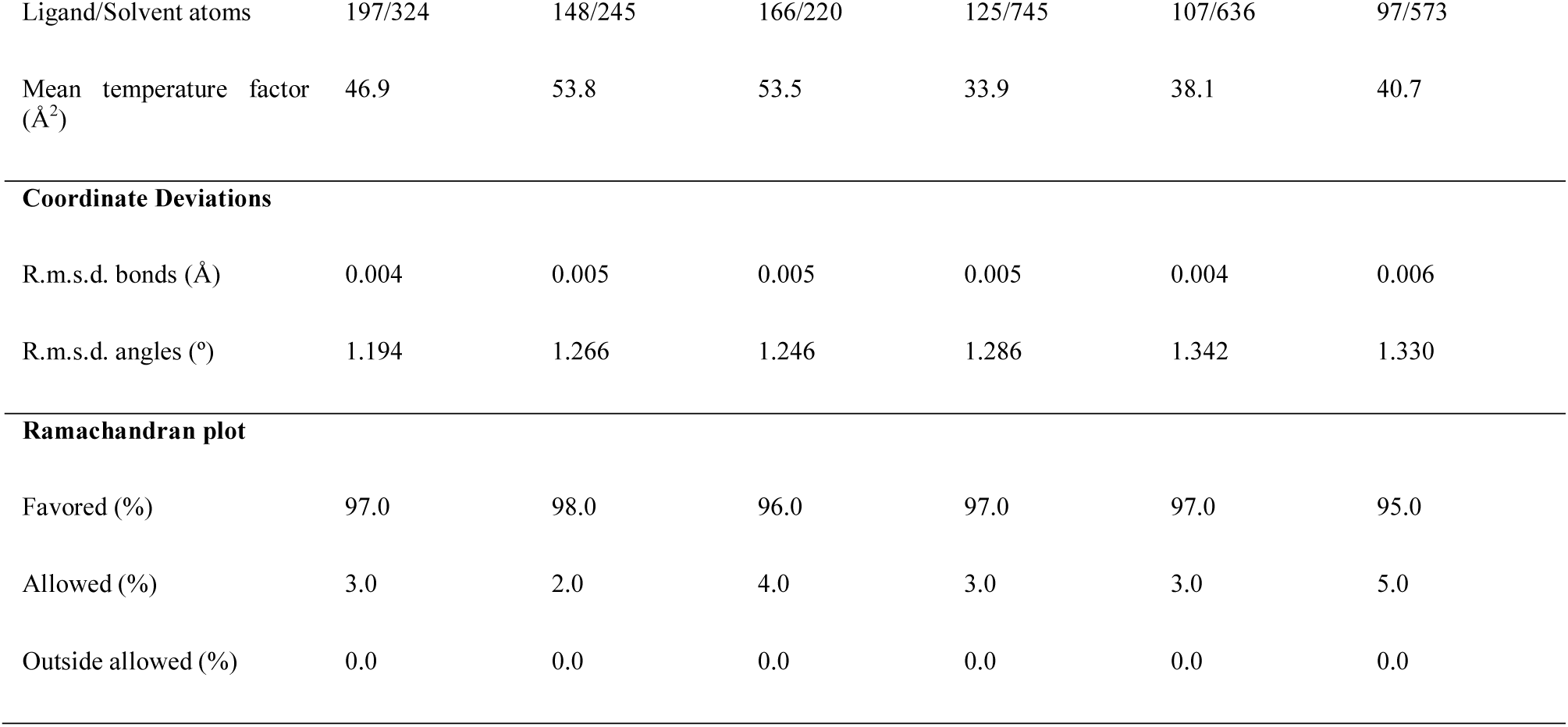
Data collection and crystallography.

| PDB Accession Code | 6WVN | 6WQ3 | 6WRZ | 6W75 | 6WKQ | 6WJT |
| --- | --- | --- | --- | --- | --- | --- |
| **Data Collection** |  |  |  |  |  |  |
| Space group | *P3<sub>1</sub>21* | *P3<sub>1</sub>21* | *P3<sub>1</sub>21* | *P3<sub>2</sub>21* | *P3<sub>2</sub>21* | *P3<sub>2</sub>21* |
| Unit cell parameters (Å; °) | *a* = *b* = 169.4,<br>*c* = 52.1;<br>*α* = *β* = 90.0,<br>*γ* = 120.0 | *a* = *b* = 169.6,<br>*c* = 52.1;<br>*α* = *β* = 90.0,<br>*γ* = 120.0 | *a* = *b* = 169.14,<br>*c* = 51.8;<br>*α* = *β* = 90.0,<br>*γ* = 120.0 | *a* = *b* = 166.2,<br>*c* = 98.3;<br>*α* = *β* = 90.0,<br>*γ* = 120.0 | *a* = *b* = 166.2,<br>*c* = 98.1;<br>*α* = *β* = 90.0,<br>*γ* = 120.0 | *a* = *b* = 166.9,<br>*c* = 98.1;<br>*α* = *β* = 90.0,<br>*γ* = 120.0 |
| Resolution range (Å) | 30.00 - 2.00<br>(2.03 - 2.00) | 30.00 - 2.10<br>(2.14 - 2.10) | 30.00 - 2.25<br>(2.29 - 2.25) | 30.00 - 1.95<br>(1.98 - 1.95) | 30.00 - 1.98<br>(2.01 - 1.98) | 30.00 - 2.00<br>(2.03 - 2.00) |
| No. of reflections | 58,029 (2,869) | 50,035 (2,464) | 40,141 (2,001) | 113,483 (5,592) | 108,598 (5,383) | 105,884 (5,270) |
| *R*<sub>merge</sub> (%) | 7.0 (86.7) | 7.5 (81.1) | 7.7 (85.7) | 7.7 (74.2) | 7.1 (78.4) | 10.1 (79.5) |
| Completeness (%) | 99.8 (100.0) | 98.8 (98.9) | 99.0 (99.6) | 100.0 (100.0) | 100.0 (100.0) | 100.0 (100.0) |
| *⟨I/σ(I)⟩* | 20.3 (2.1) | 21.1 (2.3) | 23.4 (2.8) | 25.8 (3.2) | 22.2 (2.0) | 16.7 (3.2) |
| Multiplicity | 5.1 (5.1) | 7.7 (7.8) | 7.7 (7.3) | 7.6 (7.6) | 5.5 (5.2) | 6.5 (6.6) |
| Wilson *B* factor | 37.4 | 45.4 | 45.1 | 28.4 | 31.7 | 34.2 |
| **Structure Determination** |  |  |  |  |  |  |
| MR initial model (PDB ID) | 6W4H | 6W4H | 6W4H | 6W4H | 6W75 | 6W75 |
| **Refinement** |  |  |  |  |  |  |
| Resolution range (Å) | 29.35 - 2.00<br>(2.05 - 2.00) | 28.31 - 2.10<br>(2.15 - 2.10) | 29.91 - 2.25<br>(2.31 - 2.25) | 29.93 - 1.95<br>(2.00 - 1.95) | 29.92 - 1.98<br>(2.03 - 1.98) | 29.79 - 2.00<br>(2.05 - 2.00) |
| Completeness (%) | 99.8 (100.0) | 98.8 (99.1) | 99.0 (99.4) | 99.9 (99.4) | 100.0 (100.0) | 99.8 (98.3) |
| No. of reflections | 54,988 (4,221) | 47,173 (3,646) | 38,166 (2,932) | 113,455 (8,247) | 108,429 (7,960) | 100,620 (5,598) |
| *R*<sub>work</sub>/*R*<sub>free</sub> (%) | 16.2/17.8<br>(24.9/25.1) | 16.6/18.6<br>(23.8/26.3) | 16.2/19.0<br>(21.2/21.5) | 15.7/17.5<br>(21.5/23.2) | 16.2/18.0<br>(25.1/25.4) | 17.1/19.1<br>(24.5/26.9) |
| Protein chains/atoms | 2/3,252 | 2/3,223 | 2/3,214 | 4/6,384 | 4/6,408 | 4/6,400 |
| Ligand/Solvent atoms | 197/324 | 148/245 | 166/220 | 125/745 | 107/636 | 97/573 |
| Mean temperature factor ($\text{\AA}^2$) | 46.9 | 53.8 | 53.5 | 33.9 | 38.1 | 40.7 |
| **Coordinate Deviations** |  |  |  |  |  |  |
| R.m.s.d. bonds ($\text{\AA}$) | 0.004 | 0.005 | 0.005 | 0.005 | 0.004 | 0.006 |
| R.m.s.d. angles ($^\circ$) | 1.194 | 1.266 | 1.246 | 1.286 | 1.342 | 1.330 |
| **Ramachandran plot** |  |  |  |  |  |  |
| Favored (%) | 97.0 | 98.0 | 96.0 | 97.0 | 97.0 | 95.0 |
| Allowed (%) | 3.0 | 2.0 | 4.0 | 3.0 | 3.0 | 5.0 |
| Outside allowed (%) | 0.0 | 0.0 | 0.0 | 0.0 | 0.0 | 0.0 |

### Structure solution and refinement

The first structure of nsp16/nsp10 from SARS-CoV-2 in complex with SAM with the small unit cell was determined by Molecular Replacement with Phaser (48) from the CCP4 Suite (49) using the crystal structure of the nsp16/nsp10 heterodimer from SARS-CoV as a search model (PDB ID 3R24, (14)). For all other crystal structures, refined structure from this crystal form was used as a search model. The initial solutions went through several rounds of refinement in REFMAC v. 5.8.0258 (50) and manual model corrections using Coot (51). The water molecules were generated using ARP/wARP (52), SAM, SAH or SFG, Zinc ions and ligands were added to the model manually during visual inspection in Coot. Translation–Libration–Screw (TLS) groups were created by the TLSMD server (53) (http://skuldbmsc.washington.edu/~tlsmd/) and TLS corrections were applied during the final stages of refinement. MolProbity (54) (http://molprobity.biochem.duke.edu/) was used for monitoring the quality of the model during refinement and for the final validation of the structure. A total of six structures were deposited to the Protein Data Bank (https://www.rcsb.org/) with the assigned PDB codes 6W75, 6WJT, 6WQK, 6WQ3, 6WVN, and 6WRZ with associated validation reports including electron density maps of all ligands of Interest.

### Structural alignment

The PDB coordinates of SARS-CoV nsp16 and nsp10 were analyzed on the FATCAT (29) and PDBFlex servers (55) to perform structural and sequence alignment. Structural alignments and structure figures were downloaded from the servers and modeled in Pymol open source V 2.1 (56).

## Acknowledgments

The authors thank Lukasz Jaroszewski and Adam Godzik for construct design and Grant Wiersum for protein expression. This project has been funded in whole or in part with Federal funds from the Department of Health and Human Services. National Institutes of Health, National Institute of Allergy and Infectious Diseases under Contract No. HHSN272201700060C. This research used resources of the Advanced Photon Source, a U.S. Department of Energy (DOE) Office of Science User Facility operated for the DOE Office of Science by Argonne National Laboratory under Contract No. DE-AC02-06CH11357. Use of the LS-CAT Sector 21 was supported by the Michigan Economic Development Corporation and the Michigan Technology Tri-Corridor (Grant 085P1000817).

## Author contributions

M.R.-L. and O.K. purified proteins and formed the complex, L.S. conducted crystallization and soaking experiments G.M. and J.B. collected crystallographic data, determined and analyzed structures. M.R.-L. and N.L.I wrote the draft of the manuscript, which was edited by all authors. K.J.F.S. supervised all aspects of the project.

## Data and materials availability

All data are publicly available in the RCSB Protein Data Bank (www.rcsb.org). Plasmids have been deposited and are available from www.BEIresources.com. Raw x-ray diffraction data for small unit cell and large unit cell crystals are deposited at proteindiffraction.org.

